# Identification of Rho-dependent termination site *in vivo* using synthetic sRNA

**DOI:** 10.1101/2022.09.26.509630

**Authors:** Xun Wang, Monford Paul Abishek N, Heung Jin Jeon, Jin He, Heon M. Lim

## Abstract

Rho promotes Rho-dependent termination (RDT) at the Rho-dependent terminator, producing a variable-length region at the 3′-end of mRNA without secondary structure. Determining the exact RDT site *in vivo* is challenging because the 3′-end of mRNA is rapidly removed by 3′- to 5′-exoribonuclease digestion after RDT. Here, we applied synthetic sRNA (sysRNA) to pinpoint RDT sites *in vivo* by exploiting its complementary base-pairing ability to target mRNA. Through the combined assays of rapid amplification of cDNA 3′-ends, primer extension, and capillary electrophoresis, we could precisely locate and quantify mRNA 3′-ends. We found that complementary double-stranded RNA (dsRNA) formed between sysRNA and mRNA was efficiently cleaved by RNase III in the middle of the dsRNA region. The formation of dsRNA seems to protect the cleaved RNA 3′-ends from rapid degradation by 3′- to 5′-exonuclease, thereby stabilizing the mRNA 3′-end. We further verified that the signal intensity at the 3′-end was positively correlated with amounts of mRNA. By constructing a series of sysRNAs with target sites in close proximity, and comparing the difference in signal intensity at the 3′-end of wild-type and Rho-impaired strains, we finally identified a region of increased mRNA expression within 21 bp range, which was determined as RDT site. Our results demonstrated the ability to use sysRNA as a novel tool to precisely localize RDTs *in vivo* and expanded the range of sysRNA applications.

**IMPORTANCE:** With the emergence of more new tools for inhibiting gene expression, sysRNA, which was once widely used, has gradually faded out of people′s attention due to its unstable inhibition effect and low inhibition efficiency. However, it remains an interesting topic as a regulatory tool due to its ease of design and low metabolic burden on cells. Here, for the first time, we discovered a new function to identify RDT sites *in vivo* using sysRNA. This new feature is important because, since the discovery of the Rho protein in 1969, it has been difficult to specifically identify RDT sites *in vivo* due to the rapid processing of RNA 3′-ends by exonucleases, and sysRNA might provide a new way to address this challenge.

## INTRODUCTION

Bacteria utilize transcription terminators to terminate transcription, which are classified into Rho-independent termination (RIT) and Rho-dependent termination (RDT) according to sequence properties and regulatory mechanisms (1). The Rho-independent terminator consists of two main modules: a guanine-and cytosine-rich RNA stem-loop structure and a 7-8 bp uridine-rich downstream sequence called the U-tract. When the U-tract is transcribed, it stalls RNA polymerase long enough to allow stem loop formation that disrupts the transcriptional complex, resulting in transcription termination in the U-tract (2–4). Since the sequence features required for RIT function are well defined and transcription termination actually occurs at the U-tract (5, 6), precise localization of the RIT sites *in vivo* is not difficult.

Rho is a ring-shaped homo-hexameric protein with RNA-dependent ATPase activity (7). This ATPase activity enables it to translocate along the nascent transcripts and mediate the release of RNA transcripts from halted transcription complex at the termination region (8–10). The Rho-dependent terminator also consists of two main modules: a cytosine-rich and guanine-poor Rho-binding site (C-rich region) lacking secondary structure, and a termination region downstream of the C-rich region (11). Computational prediction of Rho-dependent terminators is not straightforward due to the complex and poorly defined sequence features required by the Rho function. Nadiras et al. (2018) and Salvo et al. (2019) constructed prediction models for Rho-dependent terminators (12, 13). However, Chhakchhuak et al. (2022) experimentally confirmed that the accuracy of the model need to be improved (14). Therefore, experimental methods to identify RDT loci are highly required.

There are two main methods of detecting terminators *in vivo*, depending on where the terminator is located: one on the plasmid and the other on the chromosome. The first approach is to construct a series of plasmids, clone the terminator upstream of the reporter gene, and determine the strength of the terminator by comparing the expression level of the reporter gene (15, 16). Using this method, John et al. (1985) and Hart et al. (1991) identified the Rho-dependent terminator trp t’ of the tryptophan operon and the Rho-dependent terminator tR1 of the λ phage, respectively. In *Escherichia coli*, transcription is coupled with translation, and translating ribosomes repress RDT by sterically blocking Rho entry into nascent transcripts (17). Decoupling of translation and transcription allows Rho to bind to the C-rich region and induce transcription termination (18). The terminators of trp t’ and tR1 are located in the intergenic region and not translated. When the terminator is cloned into the plasmid, the region upstream of the reporter gene is also not translated. Thus, terminators are in an untranslated state on both plasmids and chromosomes, and they are not subject to translational repression. On the other hand, two potential Rho-dependent terminators, *tiZ1* and *tiZ2*, located in the intragenic region of *lacZ*, are inactive when the *lacZ* mRNA is translated (18, 19). When the terminator is cloned from the chromosome to the plasmid, the translation state of the terminator changes from the translated status to the untranslated state. Subsequently, the terminators changes from the inhibited state to the active state. Therefore, the above method cannot reflect the termination effect in its normal physiological condition.

The second method, *in situ* detection of Rho-dependent terminators on chromosomes, is an approach to avoid the problem of altered translation status. Researchers treated bacterial strains with antibiotic bicyclomycin (BCM) to impair Rho activity, thereby increasing transcriptional read-through at RDT sites (20). Peters et al. (2009, 2012) examined changes in the distribution of *E. coli* RNA polymerase in response to BCM by chromatin immunoprecipitation and microarrays (ChIP-chip) as well as RNA sequencing (21, 22). Sequences in transcriptional read-through regions are considered to contain RDT sites. Using this method, they identified over 1,200 RDT loci on a genome-wide scale. Later, Passong and Ranjan (2022) measured the termination efficiency of 72 Rho-dependent terminators in *E. coli* by quantitative reverse transcription PCR (RT-qPCR) analyses (14). In this approach, primers were designed downstream of suspected RDT sites, and RDT regions were validated by comparing RNA expression levels in wild-type strains and strains with impaired Rho activity. Indeed, due to the lack of secondary structure (23), the 3′-end of Rho-terminated mRNAs is rapidly digested by 3′- to 5′-exoribonuclease digestion after RDT. As a result, these identified sites are hundreds or thousands of bases long, far exceeding the length of Rho-dependent terminators. Those read-through regions in Rho-impaired strain actually consist of three parts: (1) the exonuclease-processed sequences; (2) the C-rich regions and RDT sites and (3) the read-through sequences downstream of the RDT sites. Therefore, neither methods can precisely locate the RDT sites. On top of that, Dar et al. (2018) attempted to uncover unprocessed RNA 3′-ends by reducing 3′- to 5′-exonuclease activity in *E. coli* (24). However, various exonucleases such as polynucleotide phosphorylase (PNPase), RNase II, or RNase R can process the 3′-end of RNA (25, 26). Since they could not obtain double- or triple-deletion mutants (24, 25), they were unable to solve the problem of exonuclease digestion.

sRNAs play a vital role in regulating gene expression in response to extracellular stress by base-pairing with mRNA or other sRNAs (27–30). Most sRNAs are divided into two parts: a 3′-Hfq-binding scaffold sequences and a 5′-target-binding sequence (27, 31, 32). These structural motifs have facilitated the development of artificially engineered synthetic sRNAs (sysRNAs) for gene regulation. For example, Na et al. (2013) designed sysRNAs with an 81 bp Hfq-binding MicC scaffold sequence at the 3′-part and 24 bp target-binding sequence at the 5′-part (33). These sysRNAs were designed to repress target gene expression at the translational level (33–35). We note that sysRNA differs from natural sRNA in that the sequence at the 5′-end of sysRNA is perfectly complementary to the target mRNA, whereas natural sRNA is only partly complementary. Since exonuclease activity is inhibited by double-stranded RNA (dsRNA) (36), we hoped to stabilize the 3′-end of mRNA by inhibiting exonucleases activity through sysRNA-mRNA interaction. We hypothesized that if the RNA 3′-end could be stabilized, the RDT-terminated transcripts would no longer be degraded by exonucleases, which would greatly improve the accuracy of our detection of RDT site *in vivo*.

In this study, to precisely locate RDT site *in vivo*, we designed sysRNA with an approximately 20 bp of target-binding sequence that perfectly base-pairs with the *galM* mRNA. Our results showed that dsRNA formed by sysRNA-mRNA hybridization recruited the endonuclease RNase III to form a new mRNA 3′-end in the middle of the dsRNA region, thereby protecting the cleaved upstream mRNA fragment from rapid degradation by 3′- to 5′-exonucleases and consequently stabilizing the 3′-end of mRNA. We determined that the intensity of the signal at the 3′-end of the RNA was proportional to the amount of mRNA expression. Based on the above results, we designed and constructed four sysRNAs that bind to different regions upstream and downstream of the predicted *galM* RDT region. By measuring the signal intensity of these four newly generated 3′-ends by sysRNAs, we validated the region of increased mRNA expression in the Rho-impaired strain, which was identified as the *galM* RDT site. The RDT site was precisely located within 21 bp. As far as we know, this is the most accurate localization of RDT site *in vivo*. We believe that this approach can be extended to identify other Rho-dependent terminators and is important for gaining further insight into RDT.

## RESULTS

### SysRNA construction and overall design principle

To develop a method for pinpointing the RDT, we used Rho-dependent terminator downstream of *galM* as an example. In a previous study, we reported the presence of two tandem terminators, the Rho-independent terminator and the Rho-dependent terminator, located at +7 to +30 and +31 to +108 downstream of *galM*, respectively (setting the first base downstream of the *galM* translation termination site to +1) (23). Based on the *in vitro* transcription results, RNA polymerase undergoes transcriptional pause at +111 and terminates transcription at +124 (23). *In vivo*, we could not detect the precise *galM* RDT site, but only a processed and relatively stable 3′-end signal at +28, close to the RIT stem-loop structure (23). To determine the exact RDT site *in vivo*, we constructed four MicC-scaffold sysRNAs: MicC-galM1, MicC-galM2, MicC-galM3, and MicC-galM4. These four sysRNAs bind to various regions in the predicted Rho-dependent terminator downstream of the *galM* ORF, an invisible fragment of the *pre-galETKM* (Figure 1A). Further, the MicC-scaffold sRNAs were expressed under the control of the *lac* promoter in a pBR322-derived plasmid (Figure 1B). If the sysRNA synthesized *in vivo* bound to the invisible fragment of pre-*galETKM* mRNA, it would generate a dsRNA that blocks cleavage by exonucleases, resulting in a new RNA 3′-end signal on the stabilized mRNA. Therefore, we expected that the binding of the sysRNA to the invisible pre-*galETKM* mRNA would generate a 3′-end at a specific residue in different locations. Notably, if the dsRNA region blocks the digestion of incoming 3′- to 5′-exoribonuclease, the detected 3′-end of the mRNA should be a few nucleotides downstream of the sysRNA binding site. We hypothesized that in strains with impaired Rho activity, increased mRNA expression downstream of the RDT site would result in enhanced signaling at the mRNA 3′-end. Therefore, comparing the signal intensity at the 3′-end of wild type and Rho-impaired strains can help us identify RDT sites *in vivo* (Figure 1C).

**Figure 1.**
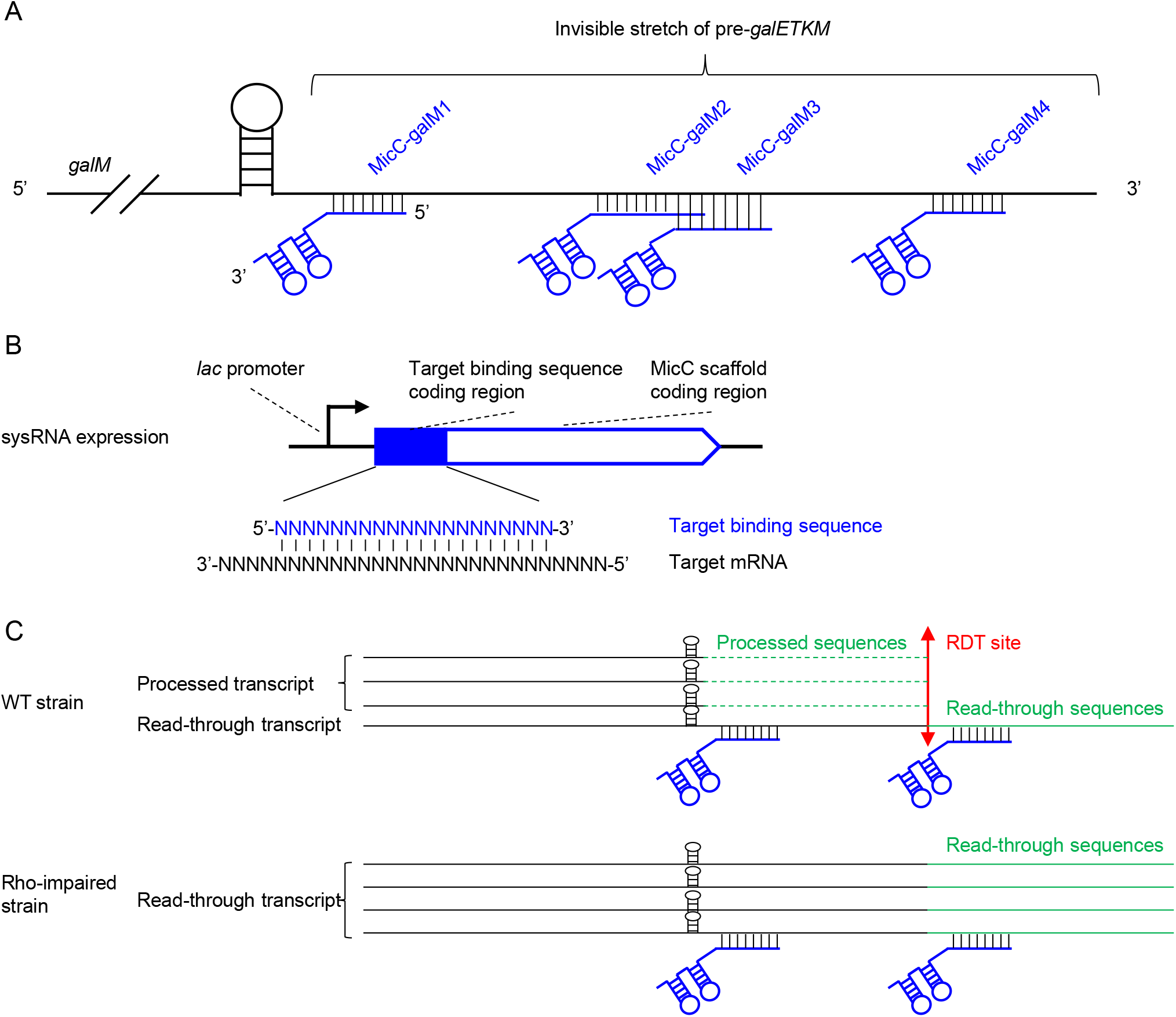
Schematic representation of overall design principles and sysRNA construction. (A) Schematic representation of the binding of the four sysRNAs on pre-*galETKM* mRNA and beyond. (B) SysRNA was expressed under the control of the *lac* promotor. The coding region of target binding sequence was inserted upstream of the MicC scaffold-coding region. (C) In the wild-type strains, RDT produces abundant terminated transcripts that are further processed by exonuclease to generate stable 3′-ends at the stem-loop structure, while the number of read-through transcripts was increased in Rho-impaired strain. The red double-arrowed line indicates the RDT site. Green dashed lines represent sequences that have been digested by exonucleases. Green straight lines represent read-through sequences.

### A new 3′-end signal was generated in the dsRNA region of the sysRNA-mRNA hybrid

To determine whether sysRNA can produce a new 3′-end signal on the invisible fragment of *pre-galETKM*, we cultivated wild-type *E. coli* MG1655 strain containing the MicC-galM expression plasmid in LB medium supplemented with ampicillin until an OD_600_ of 0.6, and extracted total RNA from the cultivated strain. The expected mRNA 3′-end was then amplified using 3′-RACE assay. Subsequently, primer extension reaction containing specific primer labeled with ^32^P was used to identify the sequence of interest in the amplified fragment. Finally, capillary electrophoresis was performed to visualize the fragment (37). In the absence of MicC-galM, no 3′-end was detected in the invisible stretch of the *pre-galETKM* mRNA (Lane 1, Figure 2A), whereas a new 3′-end was generated on the mRNA when MicC-galM was present (Lane 2-5, Figure 2A). Next, we measured the length of the DNA fragments by comparing the length of the primer extension products with that of the DNA sequencing ladders (Figure 2A). Based on the length information, we mapped the mRNA 3′-ends generated by the sysRNAs to the *gal* operon. The result showed that the sysRNAs MicC-galM1, MicC-galM2, MicC-galM3, and MicC-galM4 generated RNA 3′-ends at positions +63, +91, +111 and +155, respectively (Figure 2A). By pinpointing the 3′-ends of mRNAs, we found that they were not downstream of the binding sites of sysRNAs. Instead, all 3′-ends were formed within the dsRNA region of the sysRNA-mRNA hybrids (Figure 2B). These results suggested that, contrary to expectations, dsRNA prevented 3′- to 5′-exoribonuclease digestion to generate 3′-ends. Instead, endoribonuclease digestion resulted in the 3′-ends.

**Figure 2.**
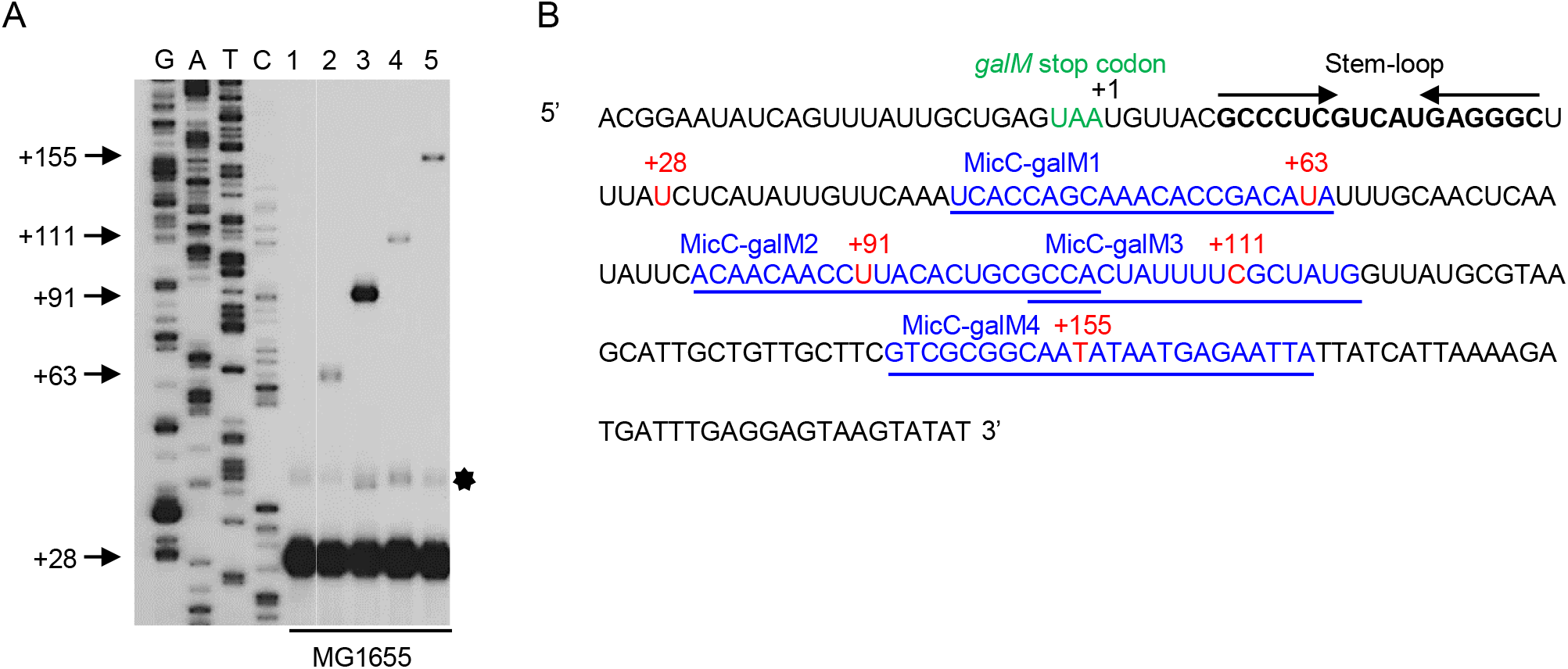
The sysRNAs enabled us to see invisible fragment of *pre-galETKM* and beyond. (A) 3′ RACE and primer extension assays of *gal* transcripts generated *in vivo* in MG1655 harboring different plasmids indicated in the figure. Lane 1, MG1655-pLac; Lane 2, MG1655-pMicC-galM1; Lane 3, MG1655-pMicC-galM2; Lane 4, MG1655-pMicC-galM3; Lane 5, MG1655-pMicC-galM4. The numbers on the left indicate the position of the 3′-ends of *gal* mRNA generated by sysRNA binding (the first base downstream of the *galM* translation termination site is set to +1). The DNA sequencing ladders in the four lanes are marked G, A, T and C, respectively. The bands indicated by the hexagon are the +28 3′ end bands, where the persistence of residual RNA secondary structures result in slower migration. (B) Sequence downstream of the *galM* stop codon. The mRNA sequences that each sysRNA can base-pair with are indicated in blue and underlined. The 3′-ends of the RNA generated by the synthetic sRNAs are shown in red. *galM* stop codon is presented in green.

### 3′-ends were generated by RNase III cleavage

To identify the endonuclease responsible for cleavage, we examined the RNA 3′-ends in *E. coli* strains that lack one of the endoribonucleases (RNase III, RNase G, RNase E, or RNase P) harboring the MicC-galM plasmid. No change in 3′-ends signal was observed in RNase III, RNase G and RNase E mutants. We found that in the RNase III control strain, all four sysRNAs generated 3′-ends (Lanes 2-5, Figure 3), whereas in the ΔRNase III strains, none of the sysRNAs generated 3′-ends (Lanes 7-10, Figure 3). Meanwhile, the only +28 3′-end of mature *galETKM* mRNA was detected from strains without sysRNA (Lane 1 and 6, Figure 3). These results indicated that the +63, +91, +111 and +155 3′-ends were generated by RNase III cleavage of the dsRNA between sysRNA and the *gal* mRNA.

**Figure 3.**
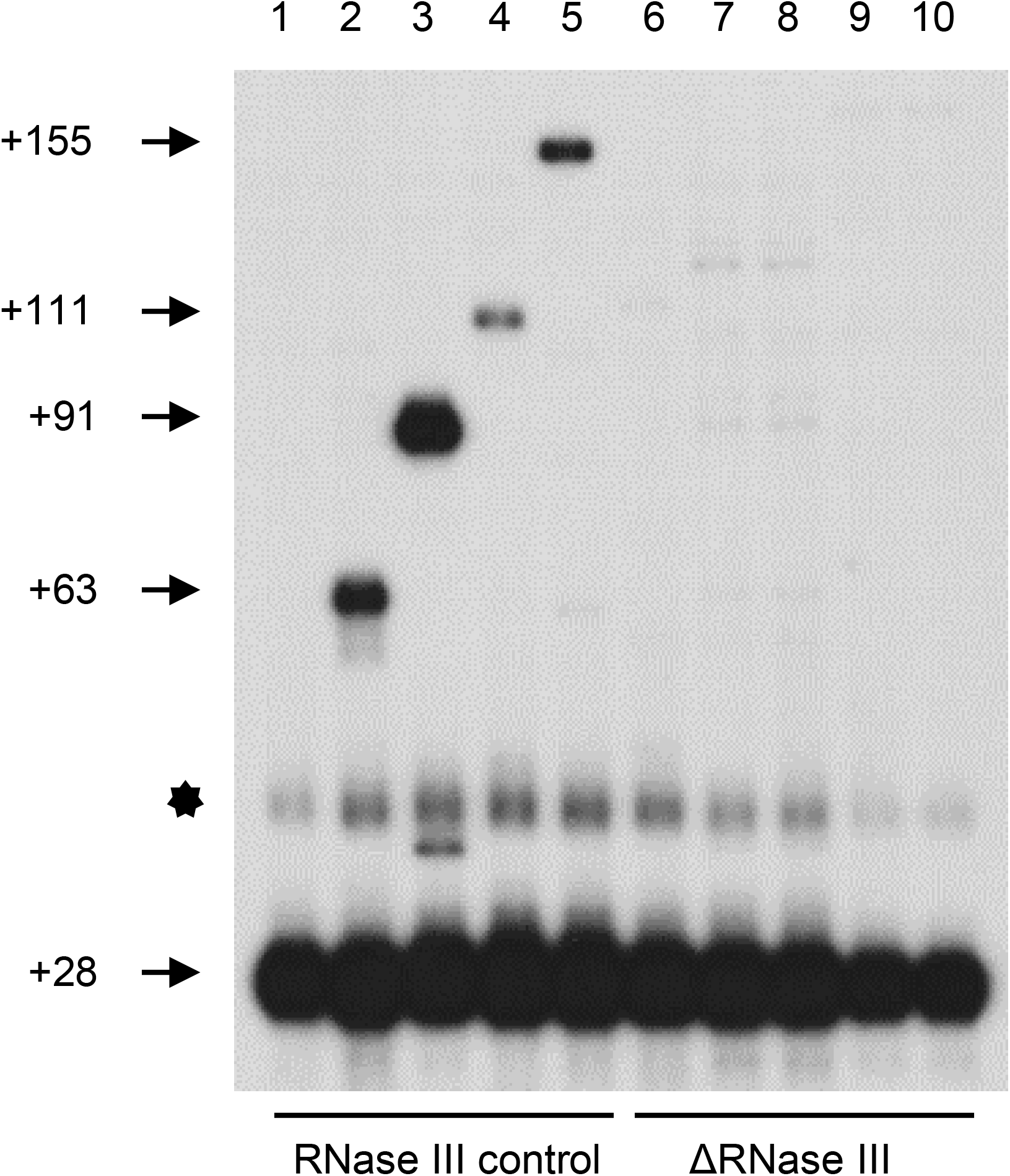
3′-RACE and primer extension assay of *gal* transcripts in RNase III control and ΔRNase III strains harboring different plasmids. Lane 1, RNase III control-pLac; Lane 2, RNase III control-pMicC-galM1; Lane 3, RNase III control-pMicC-galM2; Lane 4, RNase III control-pMicC-galM3; Lane 5, RNase III control-pMicC-galM4; Lane 6, ΔRNase III-pLac; Lane 7, ΔRNase III-pMicC-galM1; Lane 8, ΔRNase III-pMicC-galM2; Lane 9, ΔRNase III-pMicC-galM3; Lane 10, ΔRNase III-pMicC-galM4. The bands indicated by the hexagon are the +28 3′ end bands, where the persistence of residual RNA secondary structure results in slower migration.

### Amounts of mRNA and Hfq affected 3′-end signal intensity

To investigate the factors that affect the signal intensity at the 3′-end of mRNA, we checked the sysRNA expression level, mRNA expression level, and the ability of mRNA to bind sysRNA at the +91 3′-end. First, we examined the effect of the amount of MicC-galM2 on the intensity of +91 3′-end by culturing the MG1655/MicC-galM2 strain in LB medium and inducted MicC-galM2 expression with isopropyl β-D-1-thiogalactopyranoside (IPTG). We detected the expression of MicC-galM2 by dot blot assay (Figure 4A). The results showed that the expression of the MicC-galM2 increased with time, and at 8 min after IPTG induction, it was 1.5-folds that before induction (Figure 4A). Notably, MicC-galM2 exhibited relatively high expression even before induction, likely due to transcription leakage (0 min, Figure 4A). On the other hand, the expression of +91 3′-end at 0 min was the same as that at 8 min after induction (Figure 4B), suggesting that MicC-galM2 did not affect the expression of +91 3′-end when MicC-galM2 exceeded a certain threshold. Second, we investigated the effect of the amount of *gal* mRNA on the intensity of the +91 3′-end. We quantified *galM* mRNA using RT-qPCR with primers designed to amplify a region of 100 bp upstream of the *galM* stop codon UAA (Figure 4C). The results showed a 2-folds increase in *galM* mRNA expression after induction with galactose for 4 and 8 min. The expression of the +91 3′-end was also gradually increased with time, and at 8 min, it was 2.0-folds higher than before induction (Figure 4D). These results indicated that *gal* mRNA expression was positively correlated with expression at the +91 3′-end. Finally, we explored the effect of Hfq on the generation of +63, +91, +111 and +155 3′-ends. Hfq is a ubiquitous Sm-like RNA-binding protein that promotes sRNA–mRNA interactions (38, 39). We compared those 3′-ends in the MG1655 strain with those in the *Δhfq* strain harboring the pMicC-galM plasmids shown in the figure. The signal intensities at the sysRNA-generated 3′-ends in *Δhfq* by were significantly lower than in MG1655, suggesting an important role for Hfq in sysRNA function (Figure 4E).

**Figure 4.**
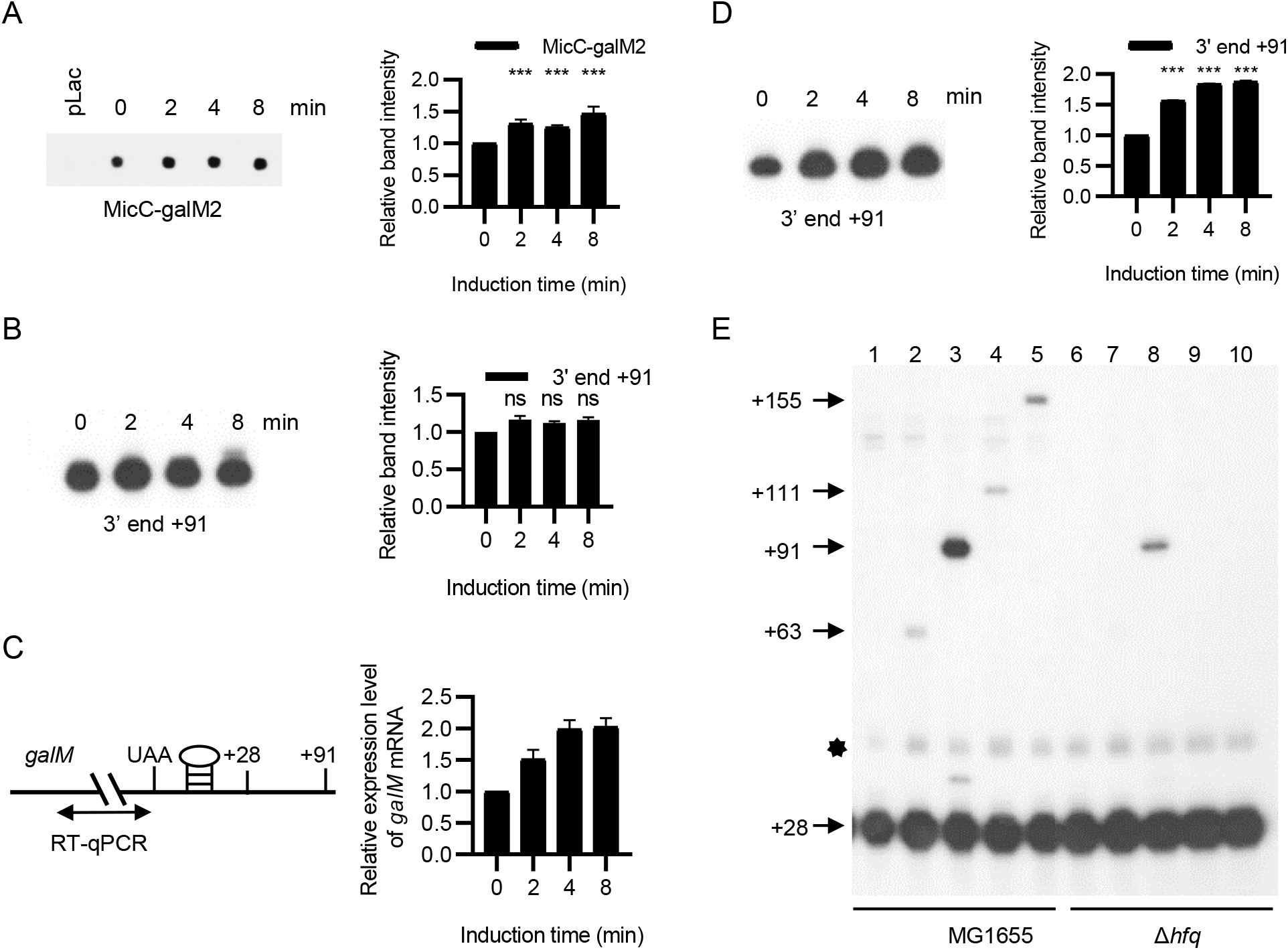
Factors affecting 3′-end generation. (A) Expression of MicC-galM2 at 0, 2, 4, and 8 min after IPTG induction. Signal intensities of each band were quantified using ImageJ. The relative expression of MicC-galM2 after induction was presented as a histogram. (B) Expression of +91 3′-end at 0, 2, 4, and 8 min after IPTG induction of MicC-galM2. Signal intensity of each band was quantified using ImageJ. The relative expression of +91 3′-end after induction was presented as a histogram. (C) Graph showing *galM* mRNA with primers (double arrow line) amplifying the region upstream of the *galM* translation stop codon. The RNA stem-loop structure indicates a Rho-independent terminator. Relative expression levels of *galM* in MG1655 strain grown in LB after galactose induction were measured by real-time quantitative PCR (RT-qPCR). (D) Expression of 3′-end +91 at 0, 2, 4, and 8 min after galactose induction. Signal intensity of each band was quantified using ImageJ. The relative expression of +91 3′-end after induction was presented as a histogram. (E) 3′ RACE and primer extension assays of *gal* mRNAs in MG1655 and *Δhfq* strains harboring different plasmids indicated in the figure. Lane 1, MG1655-pLac; Lane 2, MG1655-pMicC-galM1; Lane 3, MG1655-pMicC-galM2; Lane 4, MG1655-pMicC-galM3; Lane5,MG1655-pMicC-galM4; Lane 6, Δ*hfq*-pLac; Lane 7, Δ*hfq*-pMicC-galM1; Lane 8, Δ*hfq*-pMicC-galM2; Lane 9, Δ*hfq*-pMicC-galM3; Lane 10, Δ*hfq*-pMicC-galM4. The numbers on the left represent the location of the 3′-ends. The bands indicated by the hexagon are the +28 3′ end bands, where the persistence of residual RNA secondary structure results in slower migration. Data represent mean ± SD of 3 biological replicates.

The above results showed that the intensity of the signal at the 3′-end did not increase when sysRNA was overexpressed. However, when the amount of mRNA increased, the intensity of the signal at the 3′-end increased, and the expression of Hfq had a strong effect on the intensity of the signal at the 3′-end.

### *In vivo galM* RDT locus identification

To determine the precise RDT *in vivo*, we chose to compare the signal intensities at the +63, +91, +111 and +155 3′-ends in the Rho-impaired strain HME60 (23, 40) with those in MG1655 harboring pMicC-galM plasmids shown in the figure. We found that the signal intensities at the +63 and +91 3′-ends were the same in both strains, but those at +111 and +155 were increased by 3.5- and 2.0-folds in HME60, respectively (Figure 5A and 5B). We also found that the expression of the +28 3′-end was significantly reduced in HME60. This may be due to a decrease in Rho-terminated transcripts, resulting in a reduction in transcripts processed to +28 by exoribonucleases (23). Next, we quantified the amounts of sysRNA by dot blot and found that the expression of four sysRNAs was significantly elevated in HME60 than in MG1655 (Figure 5C and 5D). Although the expression of sysRNAs were significantly increased in HME60, the expression at the +63 and +91 3′-ends was not, confirming our previous conclusion that the increase of sysRNA could not affect the signal intensity of mRNA 3′-end when sysRNAs were in excess. Likewise, we hypothesized that the increase in signals at the +111 and +155 3′-ends was not due to increased expression of sysRNAs. We also examined the expression of *hfq* in tboth strains by RT-qPCR, and the results showed that there was no significant change in the expression of *hfq* (Figure 5E). From the above findings, we inferred that the increase at the +111 and +155 3′-end signals was due to increased mRNA expression. Therefore, *in vivo* RDT loci would be located within the +91 and +111 region.

**Figure 5.**
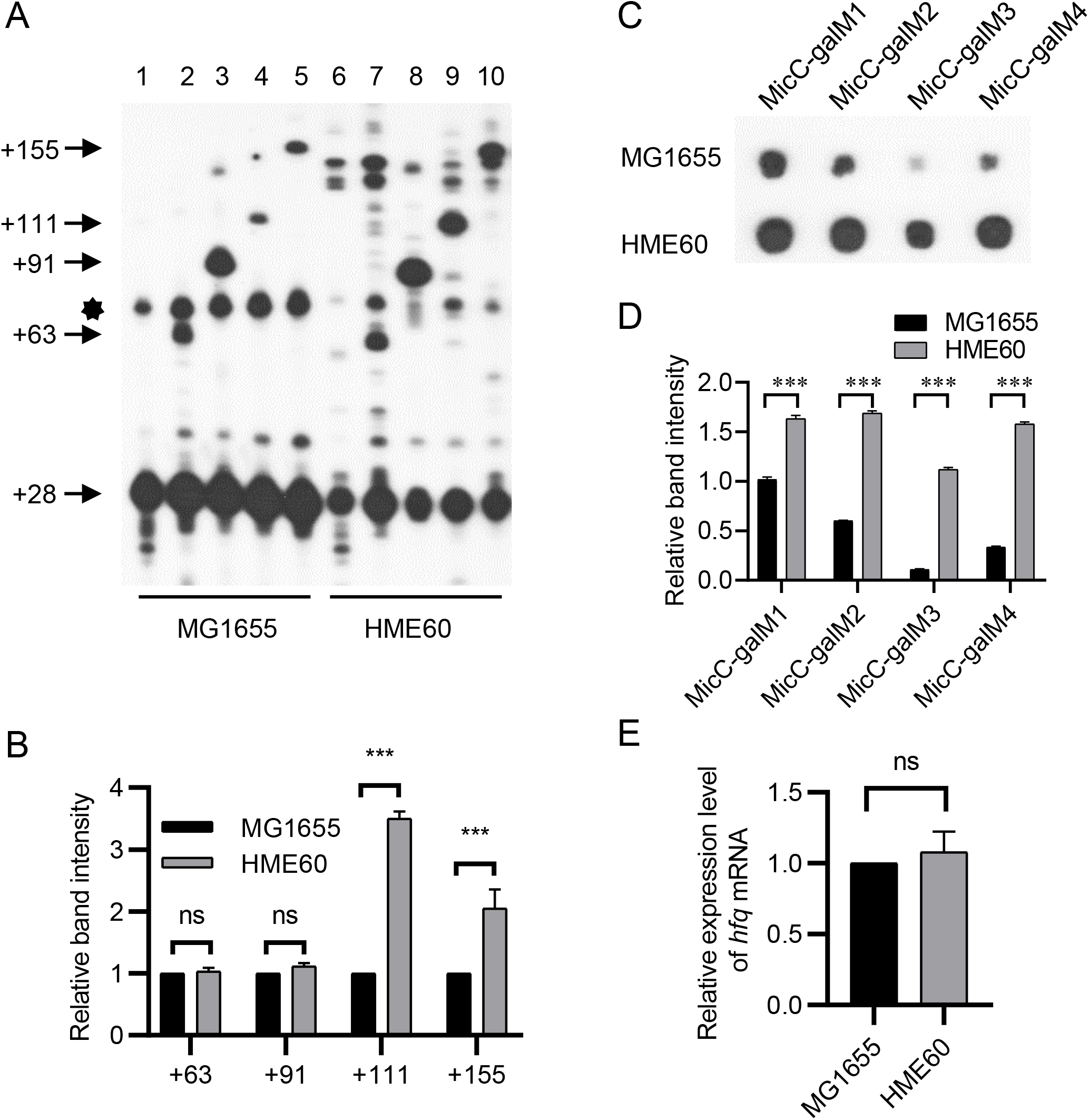
Quantification of mRNA 3′-ends between MG1655 and HME60 strains. (A) 3′-RACE and primer extension assays of *gal* transcripts in MG1655 and HME60 strains harboring different plasmids as indicated in the figure. The bands indicated by the hexagon are the +28 bands, where the persistence of residual RNA secondary structure results in slower migration. Lane 1, MG1655-pLac; Lane 2, MG1655-pMicC-galM1; Lane 3, MG1655-pMicC-galM2; Lane 4, MG1655-pMicC-galM3; Lane 5, MG1655-pMicC-galM4; Lane 6, HME60-pLac; Lane 7, HME60-pMicC-galM1; Lane 8, HME60-pMicC-galM2; Lane 9, HME60-pMicC-galM3; Lane 10, HME60-pMicC-galM4. (B) Quantification of signal intensities of 3′-ends at positions +63, +91, +111 and +155 in (A) were carried out using ImageJ. The relative density of each band is presented as a histogram. (C) Expression of four sysRNA in MG1655 and HME60 strains. Columns 1, 2, 3 and 4 show data from MicC-galM1, MicC-galM2, MicC-galM3 and MicC-galM4, respectively. (D) Quantification of signal intensity of each band was quantified using ImageJ. Relative band intensities are presented in histograms. (E) Relative expression levels of *hfq* mRNA measured by RT-qPCR in MG1655 and HME60 strains. Data represent mean ± SD of 3 biological replicates.

To further confirm our conclusions, we revalidated the RDT loci of mutated strains in the C-rich region. We first cloned the coding region from galactose operon to *gpmA* on the MG1655 genome into the single-copy plasmid pCC1BAC to generate pGal-*gpmA* plasmid. We then replaced all cytosines in the C-rich region of the *galM* Rho-dependent terminator with guanine (Figure 6A) to generate the pRDT° plasmid (23). Finally, these plasmids were individually introduced into galactose operon-deleted strain Δ*gal* for subsequent tests. We named these two derived strains as Δ*gal*-pGal-*gpmA* and Δ*gal-*pRDT°, respectively. Δ*gal*-pRDT° has been reported to be incapable of undergoing Rho-dependent transcription termination (23). The plasmid pMicC-galM was transformed into Δ*gal*-pGal-*gpmA* and Δ*gal-*pRDT° for subsequent analysis. Due to the mutations of the *gal* mRNA, MicC-galM1 and MicC-galM2 could not base pair fully complementary to the target mRNA, and we could not observe the +63 and +91 3′-ends (Figure 6B). In the Δ*gal-*pRDT° strain, we observed a 6.5- and 4.6-fold increase at the +111 and +155 3′-ends, respectively (Figure 6B and 6C). Dot blot assays of MicC-galM3 and MicC-galM4 showed that the expression of MicC-galM3 was slightly increased in the Δ*gal*-pRDT° strain, while the expression of MicC-galM4 did not change significantly (Figure 6D and 6E). Finally, we examined the expression of *hfq* in both strains by RT-qPCR and found no significant change in the expression of *hfq* in both strains (Figure 6F). Taken together, we concluded that the *galM* RDT site was located between the +91 and +111 regions.

**Figure 6.**
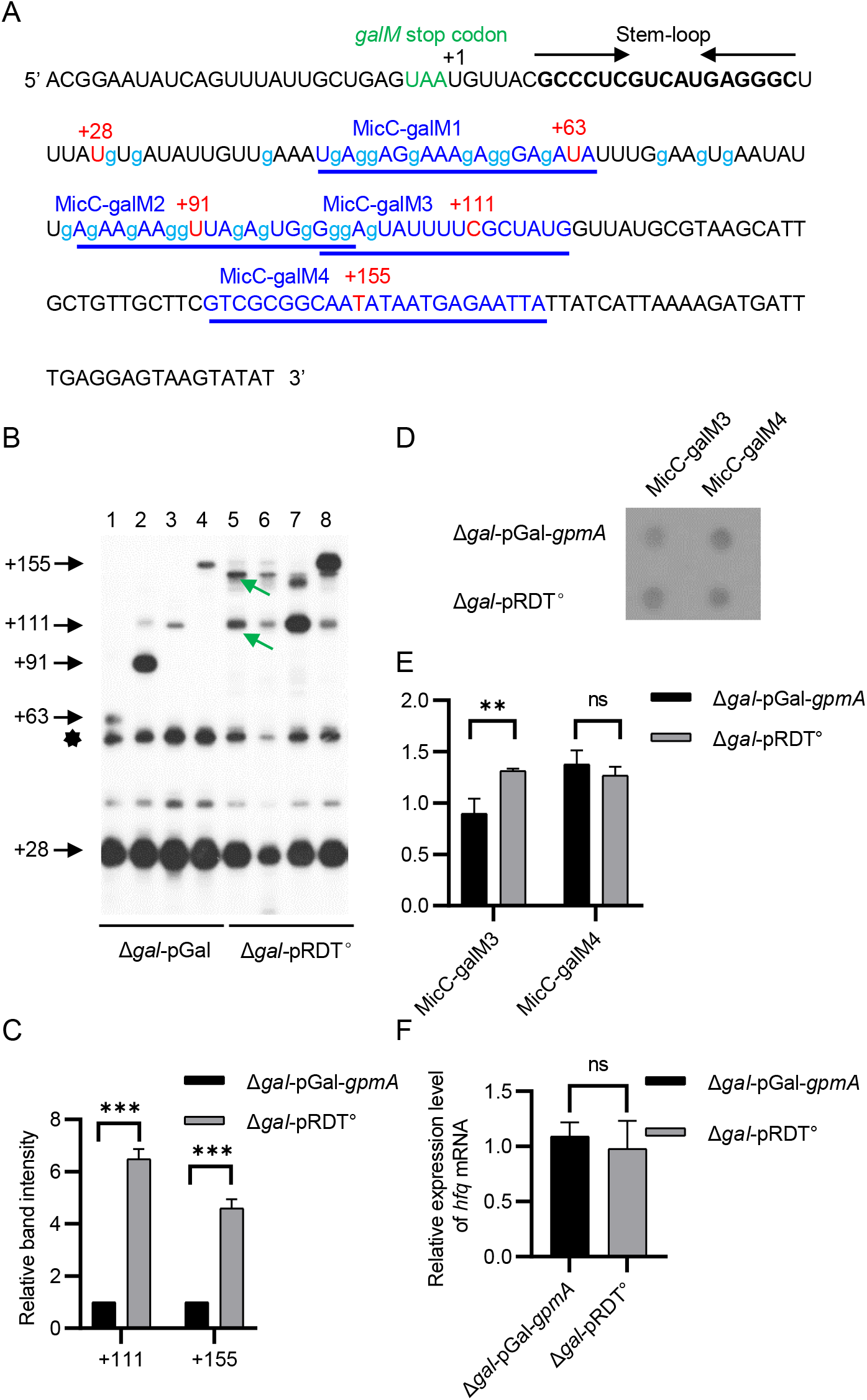
Quantification of mRNA 3′-ends between *Δgal-pGal* and Δ*gal-*pRDT°strains. (A) pRDT^°^ sequence downstream of the *galM* stop codon. Lowercase g represents mutated cytosines. Those cytosines in the pGal were mutated to guanines. (B) 3′-RACE and primer extension assays of *gal* transcripts in the Δ*gal*-pGal and Δ*gal*-pRDT° strains harboring different plasmids as indicated in the figure. The bands indicated by hexagons are the +28 bands, where the persistence of residual RNA secondary structure results in slower migration. Green arrows indicate that the new 3′-end may have arisen from changes in secondary structure due to changes in RNA sequence. Lane 1, Δ*gal*-pGal-pMicC-galM1; Lane 2, Δ*gal*-pGal-pMicC-galM2; Lane 3, Δ*gal*-pGal-pMicC-galM3; Lane 4, Δ*gal*-pGal-pMicC-galM4; Lane 5, Δ*gal*-pRDT°-pMicC-galM1; Lane 6, Δ*gal*-pRDT°-pMicC-galM2; Lane 7, Δ*gal*-pRDT°-pMicC-galM3; Lane 8, Δ*gal*-pRDT°-pMicC-galM4. (C) Signal intensities of +111 and +155 3′-ends in (B) were quantified using ImageJ. The relative density of each band is presented as a histogram. (D) Expression of MicC-galM3 and MicC-galM4 in *Δgal-pGal* and Δ*gal*-pRDT° strains. Columns 1 and 2 show MicC-galM3 and MicC-galM4, respectively. (E) The signal intensity of each band was quantified using ImageJ. Relative band intensities are presented as a histogram. (F) Relative expression levels of *hfq* mRNA measured in Δ*gal*-pGal and Δ*gal-*pRDT° strains. Data represent mean ± SD of 3 biological replicates.

### SysRNA targeting sites selection

As shown in our results, each MicC-galM generated a different amount of RNA 3′-end. The amount of +91 3′-end was much higher than the amount of +63 3′-end Figure 2). However, dot blot quantification of each MicC-galM showed no correspondence between the amount of MicC-galM RNA and the 3′-end (Figure 5C). To clarify the reasons for the difference in 3′-end generation, we analyzed the possible reasons from the perspective of *gal* mRNA. First, in terms of mRNA quantity, the two MicC-galM1 and MicC-galM2 targeting sites were within the C-rich region of *galM* Rho-dependent terminator upstream of the RDT site. Thus, the two sysRNAs would bind exactly the same amount of *gal* mRNA in this narrow region. Next, we analyzed the secondary structure of *gal* mRNA. Using *RNAfold* WebServer (48), we investigated the secondary structure of this mRNA extending from +1 to +117 at the 3′-end (Figure 7A). *RNAfold* WebServer predicted the formation of a 5 bp stem with a 10 bp loop and a 3 bp stem with a 10 bp loop in the regions of +50-+69 and +98-+113, respectively. In addition, no other secondary structure was observed within the mRNA segment. The minimum free energy (MFE) of these two stem-loops were −0.90 kcal/mol (5 bp stem with 10 bp loop) and −1.00 kcal/mol (3 bp stem with 10 bp loop). The target-binding regions of the MicC-galM1 and MicC-galM2 in the fragment showed that 15 (out of 21) and 6 (out of 22) bp sequences resided in these two secondary structures, respectively (Figure 7A). Thus, approximately 71 % of MicC-galM1-targeted mRNA sequence and 27 % of MicC-galM2-targeted mRNA sequence were located in the RNA secondary structures, suggesting that MicC-galM2 could bind target mRNA better than MicC-galM1. The relative RNase III cleavage efficiency on dsRNA was in good agreement with the percentage of base pairings in the RNA secondary structures, namely, the RNase III cleavage efficiency on dsRNA generated by MicC-galM2 was higher than that of MicC-galM2. To further reveal the effect of secondary structure on cleavage efficiency, we selected the *galM* RIT region containing robust secondary structure with a MFE of −10.30 kcal/mol as the target of the new sysRNA MicC-galM0. As expected, MicC-galM0 failed to produce a new 3′-end (Figure 7B). This can be explained by the inhibition of sysRNA function by secondary structure on target mRNA. Thus, the secondary structure of target mRNA should be considered when designing the target binding sequence to ensure that the sysRNA binds target mRNA in a region without secondary structure.

**Figure 7.**
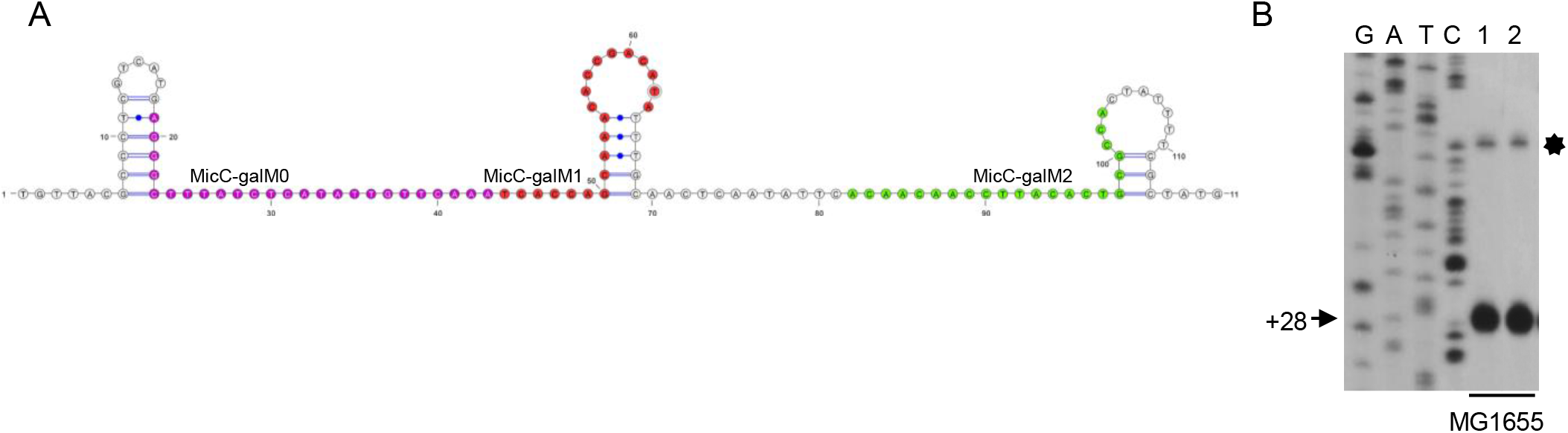
mRNA secondary structure analysis and 3′-end visualizations. (A) Secondary structure prediction of *galM* mRNA extending from +1 to +117 at the 3′-end. Strong stem-loop structure in the Rho-independent terminator of *galM* and two weak stem-loop structures in the C-rich region. Target-binding sequence of MicC-galM0 (purple), MicC-galM1 (red) and MicC-galM2 (green). (B) 3′-RACE and primer extension assays on *gal* transcripts in MG1655 strains harboring different plasmids as indicated in the figure. Lane 1, MG1655-pLac; Lane 2, MG1655-pMicC-galM0. The DNA sequencing ladders in the four lanes are labeled G, A, T and C, respectively. The bands indicated by the hexagon is the +28 band, where the persistence of residual RNA secondary structure results in slower migration.

We also designed several sysRNAs with target regions at the end of *galE* gene and the beginning of *galT* gene, and found that none of them produced the expected 3′-end. This might be attributed to the more extensive translation events in the above-mentioned regions than in other regions of the operon. Therefore, occupancy of mRNA target regions by translating ribosomes should also be considered in future studies (49–51).

## Discussion

In this study, we demonstrated that dsRNA produced by sysRNA binding to mRNA recruited RNase III for dsRNA cleavage and that the cleaved RNA 3′-end is resistant to exoribonuclease digestion, thereby stabilizing the rapidly degraded RNA 3′-end. By constructing four sysRNAs that bind to different sites of mRNA, we observed four distinct stable RNA 3′-ends. When the RNA 3′-end was upstream of the RDT site, the signal intensity at the 3′-end of MG1655 was unchanged compared to the Rho-impaired strain. However, when the RNA 3′-end was located downstream of the RDT site, the signal intensity at the RNA 3′-end increased due to the increased expression of downstream mRNA in Rho-impaired strain. By this method, we identified the RDT site of the last gene of the galactose operon *galM in vivo*. We believe that using this new tool to identify the exact RDT site could deepen our understanding of RDT.

### Possible reasons for inconsistency of RDT loci *in vitro* and *in vivo*

We note that the *galM* RDT site we detected differed between +91 and +111 *in vivo* and +124 *in vitro*. We speculated that these results are inconsistent for the following reasons: First, most RDTs are regulated by factors other than Rho proteins, including trans-acting elements such as translating ribosomes, RNA-binding proteins, Rho-binding proteins, or cis-acting elements, such as RNA-stem loops, riboswitches, etc. (41–43). Changes of these factors in the *in vivo* and *in vitro* environments can lead to differences in RDT. Second, forward transcription rates and pause sites of RNA polymerase may vary (44–47). Since RNAP pause is a crucial step in the RDT process, differences in transcriptional pause sites may also lead to differences in RDT sites. In fact, due to the inability to precisely locate transcriptional termination sites *in vivo*. no case comparing *in vivo* and *in vitro* RDT sites has been reported. It is for these reasons that it is important to develop a method that can detect RDT *in vivo* without altering its physiological environment. Of these, sysRNA-based strategy has advantages in: (1) high specificity; (2) ease of design and implementation, and (3) minimal impact on cell growth. Therefore, our method provided an excellent strategy for detecting RDT sites *in vivo*.

### Further expansion of sysRNA applications

Despite the diverse physiological functions of natural sRNAs, the application of sysRNAs is still limited to inhibiting gene expression. Here, we extended their applications by using sysRNAs for the detection of RDT loci for the first time. In eukaryotes, RNA editing is an area of great interest. Compared with DNA editing, RNA editing does not require permanent changes to the genome, and this reversible and easily regulated editing approach may have advantages in terms of safety. A major problem with editing technologies that rely on the CRISPR/Cas system is that they are dependent on the expression of exogenous editing enzymes or effector proteins. This can cause problems such as cytotoxicity or delivery difficulties (52, 53). Wei′s group discovered for the first time that a specially designed RNA has great potential for RNA editing by recruiting endogenous adenosine deaminase of RNA (ADAR) protein to produce efficient and precise editing of specific adenosine on target gene transcripts without introducing any exogenous effector protein (54, 55). According to our results, sysRNA could cleave the target mRNA by recruiting endogenous RNase III. This shared some similarities with ADAR-recruiting RNA. We believe that by appropriately modifying RNase III, multiple functions of sysRNA-based strategy such as RNA editing in bacteria could be achieved.

## MATERIAL AND METHODS

### Bacterial strains and growth conditions

*E. coli* strains MG1655, *Δgal, Δhfq* and HME60 (23, 56) (Table S1) containing the corresponding plasmids indicated in the figures were grown at 37 °C in LB medium (containing 10 g tryptone, 5 g yeast extract, and 10 g NaCl per liter of water) supplemented with 0.5% (wt/vol) galactose, chloramphenicol (15 μg/mL), ampicillin (100 μg/mL), or IPTG (1 mM) as necessary. Due to leaky expression of MicC-galMs, we did not add IPTG to the medium except for IPTG induction experiments. Endonuclease deficiency and their parent strains, including SDF204 *(rnc+;* RNase III control), SDF205 *(rnc-;* ΔRNase III), GW10 *(rng+, rne+;* control for RNase G and RNase E), GW11 *(rng::cat;* ΔRNase G), GW20 (*ams1^ts^;* RNase E temperature-sensitive mutant), NHY312 *(rnpA+;* control for RNase P), and NHY322 (*rnpA49*; RNase P temperature-sensitive mutant) were generous gifts from Dr. Y. H. Lee (KAIST, South Korea) (56) (Table S1). These strains containing the corresponding plasmids were grown in LB medium supplemented with ampicillin (100 μg/mL) at appropriate temperatures (30 °C, 37 °C, and 44 °C).

### Plasmid construction

The pGal plasmid is a derivative of pCC1BAC (Epicenter Biotechnologies, USA) and contains a coding region from *galETKM* to *gpmA* for *gal* mRNA expression. On the pRDT^°^ plasmid, all cytosines in the *galM* RDT were replaced by guanine (23). To construct the sysRNA expression plasmids (pMicC-galM1, pMicC-galM2, pMicC-galM3 and pMicC-galM4), the 81 bp MicC scaffold DNA fragment of MicC was first amplified by PCR using *E. coli* genomic DNA as a template, and the obtained PCR products were digested with *Eco*RI and *HindIII* and inserted into pLac (pBR322-derived plasmid) (57) to obtain the pHL1722 plasmid. Subsequently, 20 bp sequences complementary to +44-+64, +82-+102, +100-+117 and +145-+168 *gal* operon regions were inserted into *Aat*II and *Eco*RI of pHL1722 (Figure S1) to obtain sysRNA expression plasmids. The plasmids used were listed in Table S2.

### Total RNA preparation and 3′ RACE assay

Total RNA was extracted from 2 mL *E. coli* strains as previously described (58). For the 3′-RACE assay, total RNA (10 μg) was ligated with 2 nM synthetic RNA oligomer (Table S3) with 5′-phosphate and 3′-inverted deoxythymidine (Integrated Dna Technologies, USA). RNA ligation was performed by 10 units of T4 RNA ligase (Thermo Fisher Scientific, USA) in a volume of 20 μL for 3 h at 37 °C, supplemented with 20 units of RNasin® Ribonuclease inhibitor (Promega, USA) to prevent RNase activity. The ligated RNA was purified using a G-50 column (GE Healthcare, USA) and 1 μg of RNA was reverse transcribed in a 20 μL reaction volume at 37°C for 2 h. The reverse transcription reaction contains four units of Ominiscript™ reverse transcriptase (QIAGEN, Germany), 0.5 mM dNTP mixture, 0.4 μM of 3RP primer complementary to RNA oligomer (Table S3) and 10 units of RNasin. Finally, a 2 μL sample of this reaction was used as a PCR template. A 150 bp galactose operon-specific fragment containing the region where the predicted RNA 3′-end is located was amplified using HotStarTaq™ DNA polymerase (QIAGEN, Germany) with *gal*-specific primer M3-F and 3RP primer (Table S3). To visualize mRNA 3′-ends, PCR product was purified, and used as a template for primer extension reactions. Primer extension reaction was performed with one unit of Taq polymerase (QIAGEN, Germany), 0.2 mM dNTP mixture, 0.3 μL ^32^P-labeled *gal* specific primer M4-F (Table S3) and 2 μL PCR product in a volume of 20 μL. The ^32^P-labeled M4-F was placed inside the PCR product. Then, the primer was allowed to anneal to the DNA template and Taq polymerase was used to synthesize DNA from the DNA template until it reaches the end of the DNA. Reaction products were resolved in an 8% polyacrylamide/urea sequencing gel, and quantified with a PhosphorImager (Amersham Biosciences Crop, England).

### RNA dot blot

For RNA dot blot, we first extracted total RNA, with 10 μg of total RNA treated with DNase I (Takara, Japan) and purified by G-50 column (GE Healthcare, USA). One μg of RNA was then mixed with 2×RNA loading buffer and incubated at 80 °C for 5 min to eliminate RNA secondary structures and DNase I activity. RNA was spotted onto a positively charged nylon membrane (Thermo Fisher Scientific, USA). The membrane was adhered to a Whatman filter paper and baked at 80 °C for 1 h. MicC-probe (Table S3) was labeled with ^32^P, mixed with ULTRAhyb Ultrasensitive Hybridization buffer (Thermo Fisher Scientific, USA), and hybridized to RNA extract on nylon membrane overnight at 42 °C according to the manufacturers′ instruction (31, 58). After that, the membrane was washed twice with 2×SSC containing 0.1% SDS for 5 min at 25 °C, and then washed twice with 0.2×SSC containing 0.1% SDS for 15 min at 42 °C. Radioactive dots on the membrane were visualized by exposing to X-ray film. The signal intensities of the dots were quantified using the software ImageJ. Data were subjected to one-way analysis of variance (ANOVA) using the Bonferroni test, n = 3.

### Real time quantitative PCR (RT-qPCR)

For RT-qPCR experiments, 2 mL samples from *E. coli* strains were collected. Total RNA was extracted, and RT-qPCR was conducted essentially as previously described (40). Results for each strain were normalized to those of the *rrsB* gene coding 16S rRNA. For data analysis, technical and biological triplet data were obtained. Data were subjected to one-way analysis of variance (ANOVA) using the Bonferroni test, n = 3.

## SUPPLEMENTAL MATERIAL

Supplemental material is available online only.

FIG S1

TABLE S1

TABLE S2

TABLE S3

## ACKNOWLEDGEMENTS

This work was supported by a Basic Science Research Program grant (2020R1A2C200633611 to H.M.L.) from the National Research Foundation of Korea. This work was also supported by the National Natural Science Foundation of China (31971339 and 32171422). And was also supported by the Fundamental Research Funds for the Central Universities (2662022SKYJ004).

## AUTHOR CONTRIBUTIONS

H. M.L. conceptualized and conceived the project. X.W. and M.P.A.N. performed the experiments. X.W., M.P.A.N., H.J.J. and H. J. analyzed the data. H.M.L. and X.W. wrote the manuscript, and all authors discussed the results and commented on the manuscript.

